# Effective population size for culturally evolving traits

**DOI:** 10.1101/2021.09.09.459561

**Authors:** Dominik Deffner, Anne Kandler, Laurel Fogarty

## Abstract

Population size has long been considered an important driver of cultural diversity and complexity. Results from population genetics, however, demonstrate that in populations with complex demographic structure or mode of inheritance, it is not the census population size, *N*, but the effective size of a population, *N*_*e*_, that determines important evolutionary parameters. Here, we examine the concept of effective population size for traits that evolve culturally, through processes of innovation and social learning. We use mathematical and computational modeling approaches to investigate how cultural *N*_*e*_ and levels of diversity depend on (1) the way traits are learned, (2) population connectedness, and (3) social network structure. We show that one-to-many and frequency-dependent transmission can temporally or permanently lower effective population size compared to census numbers. We caution that migration and cultural exchange can have counter-intuitive effects on *N*_*e*_. Network density in random networks leaves *N*_*e*_ unchanged, scale-free networks tend to decrease and small-world networks tend to increase *N*_*e*_ compared to census numbers. For one-to-many transmission and different network structures, effective size and cultural diversity are closely associated. For connectedness, however, even small amounts of migration and cultural exchange result in high diversity independently of *N*_*e*_. Our results highlight the importance of carefully defining effective population size for cultural systems and show that inferring *N*_*e*_ requires detailed knowledge about underlying cultural and demographic processes.

**AUTHOR SUMMARY:** Human populations show immense cultural diversity and researchers have regarded population size as an important driver of cultural variation and complexity. Our approach is based on cultural evolutionary theory which applies ideas about evolution to understand how cultural traits change over time. We employ insights from population genetics about the “effective” size of a population (i.e. the size that matters for important evolutionary outcomes) to understand how and when larger populations can be expected to be more culturally diverse. Specifically, we provide a formal derivation for cultural effective population size and use mathematical and computational models to study how effective size and cultural diversity depend on (1) the way culture is transmitted, (2) levels of migration and cultural exchange, as well as (3) social network structure. Our results highlight the importance of effective sizes for cultural evolution and provide heuristics for empirical researchers to decide when census numbers could be used as proxies for the theoretically relevant effective numbers and when they should not.

## 1. Introduction

Cultural evolutionary dynamics are governed by individual-level cognitive processes and demographic properties of the population [Cavalli-Sforza and Feldman, 1981, Boyd and Richerson, 1985]. Archaeologists and anthropologists have been particularly interested in the ways population size might shape cultural processes [see Derex and Mesoudi, 2020, Strassberg and Creanza, 2021, for recent reviews]. When researchers consider the impact of population size on cultural evolution, they predominantly refer to the number of individuals in a population. This census population size *N* is readily observable in real-world situations and can be quantified by counting how many people are present at a certain place and time. Results from population genetics, however, have long demonstrated that in most real-world populations it is not this census size, but the effective size, *N*_*e*_, that is the correct measure to use when calculating important evolutionary parameters such as genetic diversity and divergence times between populations [Ewens, 2012].

### 1.1. What is effective population size and why do we need it?

The effective population size is a theoretical construct that links complex populations to simpler, idealised models. This way, the effective size makes it possible to directly compare any number of complex populations—each with their own complicating factors—in a way that would otherwise be impossible. A commonly used simplified model in population genetics is the Wright-Fisher model [Fisher, 1923, Wright, 1931, Ewens, 2012], and much of what we understand about evolution comes from our understanding of evolution in such idealised models. The effective population size is defined in relation to this model as the size of an ideal Wright-Fisher population that experiences genetic drift at the same rate as a particular study population (see section 2 for details).

To understand what we gain from the effective size, even if we are not particularly interested in Wright-Fisher models, let us assume there are two populations, A and B, that produce a particular cultural trait with many possible variants. We now want to know whether population size affects the number of different traits in a population. Population A has a large census population size of 1000 individuals, population B has a smaller census size of just 500. Using a theoretical model of a cultural evolutionary process [e.g. Shennan, 2001, Henrich, 2004, Powell et al., 2009, Fogarty et al., 2017], we conclude that larger populations should have larger or more complex cultural repertoires. Can we expect to find this demographic relationship in data on census population sizes and cultural repertoire sizes from both populations [e.g. Oswalt, 1976]? The answer is that—regardless of how good the model is—the relationship is unlikely to be found unless these real populations are identical in some evolutionarily important ways. If they do not have the same age structure, demographic history, or, as we show below, cultural transmission mechanisms and interaction patterns, the populations are not directly comparable, except through their relation to a simpler model—through their effective population sizes.

Imagine we now discover that, 10 generations ago, population A had a population bottleneck where its census size fell to only 10 individuals before recovering to its current value of 1000. Genetic evolution will be affected by this bottleneck for a number of generations (culture might recover from such events much faster than genetics [Fogarty and Kandler, 2020]). Both populations are otherwise identical and conform to the assumptions of the Wright-Fisher model, which we detail below. Accordingly, the effective population size of the small, stable population B is 500, the same as its census size. The effective size of population A, however, is only around 92 (see appendix 1 for calculation). We can now use results from population genetics to calculate how many cultural traits we expect to see in each population, given certain transmission mechanisms and innovation rates. For population B with *N*_*e*_ = 500, the expected number of traits is 223. For population A with *N*_*e*_ ≈ 92, we expect to see on average 41 traits in a given generation (see appendix 1 for full details). Thus, although a relationship exists between effective population size and cultural diversity, a straightforward relationship does not exist between census size and diversity. Using census numbers or more informal definitions of effective size will produce incorrect results. As real populations differ from one another and from the assumptions of an ideal Wright-Fisher model in numerous evolutionarily important ways, the ability to unify and compare them is invaluable. This is relevant for any question where we need to understand the effects of drift and selection in cultural processes across populations.

For many animal and plant species, researchers have investigated the relationship between observed census size *N* and calculated (genetic) effective size *N*_*e*_. In one large-scale study, ratios of *N*_*e*_ to *N* were found to vary between 0.19 in a species of pine tree to 3.69 in a species of mosquito [Waples et al., 2013]. This demonstrates that, across species, a large range of relationships between census and effective sizes are possible and *N*_*e*_ can also exceed the census size *N*—a possibility we expand on below. Estimates of this ratio for humans suggest that the genetic effective size is considerably lower than our census size with an *N*_*e*_ of around 10,000 compared to a census size of around 7 billion at the time of publication [Tenesa et al., 2007], possibly reflecting population bottlenecks in the past. All of this strongly suggests that in order to understand how demography affects cultural evolution, and which empirical comparisons are meaningful, we need to gain a better understanding of the cultural equivalent of *N*_*e*_.

### 1.2. Cultural effective population size - history and outline

The importance of effective population size has been partially acknowledged in the cultural evolution literature and the concept is often invoked. Henrich et al. [2016], for example, write: “The theory explicitly predicts that it is the size of the population that shares information—the effective cultural population size—that matters, and if there is extensive contact between local or linguistic groups, there is no reason to expect census population size to correspond to the theoretically relevant population”. More recently, Derex and Mesoudi [2020] also claim that effective population size “depends on both population size and interconnectedness”. However, the definition of *N*_*e*_ as “the population that shares information” is not always correct and corresponds more closely to the ‘breeding population’ rather than the effective size. We hope to show that it is not only the number of individuals sharing information, also called “cultural equivalent *N*” [Cavalli-Sforza and Feldman, 1981], but the exact details of how information is passed on between individuals that should be expected to influence cultural effective population size. Furthermore, it remains unclear whether and how different forms of population interconnectedness as well as social network characteristics might influence effective population size for culture, though intuition suggests that this influence may be strong. In a first formal treatment for cultural evolution, Premo [2016] used simulation models to investigate effective population size and its relationship with cultural complexity. In the context of models by Shennan [2001] and Henrich [2004], the results show that natural and cultural selection can weaken the relationship between census population size, cultural diversity and mean skill level. This work did not formally derive an appropriate measure for cultural effective size and examined the influence of selection within the scope of two domain-specific models.

Here we aim to provide this formal derivation and systematically examine the concept of effective population size for cultural evolution. We first introduce drift and effective population size as employed in standard models of population genetics. After deriving appropriate formulations of *N*_*e*_, we use different modeling approaches to investigate how cultural *N*_*e*_ depends on (1) the way traits are learned, namely one-to-many and frequency-dependent (i.e conformist and anti-conformist) transmission, (2) population connectedness through either migration or cultural exchange, and (3) social network structure. In each case, we relate effective numbers to the emerging levels of cultural diversity. We conclude by discussing implications for the role of demography in cultural evolution and provide heuristics for empirical researchers to decide when census numbers might be used as proxies for the theoretically relevant effective numbers.

## 2. Drift and effective population size in population genetics

We first provide a basic introduction to the Wright-Fisher population, genetic drift, and the concept of effective population size as developed for genetic evolution.

### 2.1. The Wright-Fisher population and genetic drift

The classic Wright-Fisher population is a closed, randomly mating population of *N* individuals without selection and mutation [Fisher, 1923, Wright, 1931, Fisher, 1931, Ewens, 2012]. For each discrete generation, new individuals are formed by random sampling, with replacement, of gametes produced by the previous generation. The number of offspring for a given member of the parental generation is a binomially distributed random variable with both mean and variance of approximately 1 (for haploid populations; see appendix 2 for explanation). Genetic drift describes the random change in allele frequencies from generation to generation by the chance success of some alleles relative to others [Wright, 1929, Masel, 2011, Whitlock and Phillips, 2014]. While per definition unpredictable in any particular instance, on average, drift causes populations to change in broadly systematic ways: drift reduces the number of alleles in a population, increases the differences among populations and results in higher variability of allele frequencies over time. Crucially, the magnitude of allele frequency changes due to genetic drift is inversely related to the size of the Wright-Fisher population—the larger the number of individuals, the smaller the effects of genetic drift. These consequences are illustrated in Fig. 1 which shows the frequency of one allele in populations of different sizes, *N* = 10 (Fig. 1A), *N* = 100 (Fig. 1B), *N* = 1000 (Fig. 1C) or *N* = 10000 (Fig. 1D). Colored lines show trajectories for 8 separate populations evolving over 100 generations. In small populations, random sampling of alleles leads to strong fluctuations in allele frequencies over time. After relatively few generations, populations diverge and the allele either goes to fixation or extinct. Both outcomes are expected to happen half of the time as the allele was at 50% initial frequency. The larger the population, the smaller are temporal fluctuations in allele frequency and the longer it takes until populations diverge. Given enough time, in the absence of other evolutionary forces, even very large populations will diverge as much as smaller populations and the allele will go to fixation/extinction. Population size, thus, affects the rate of drift but not its eventual outcome.

**Figure 1.**
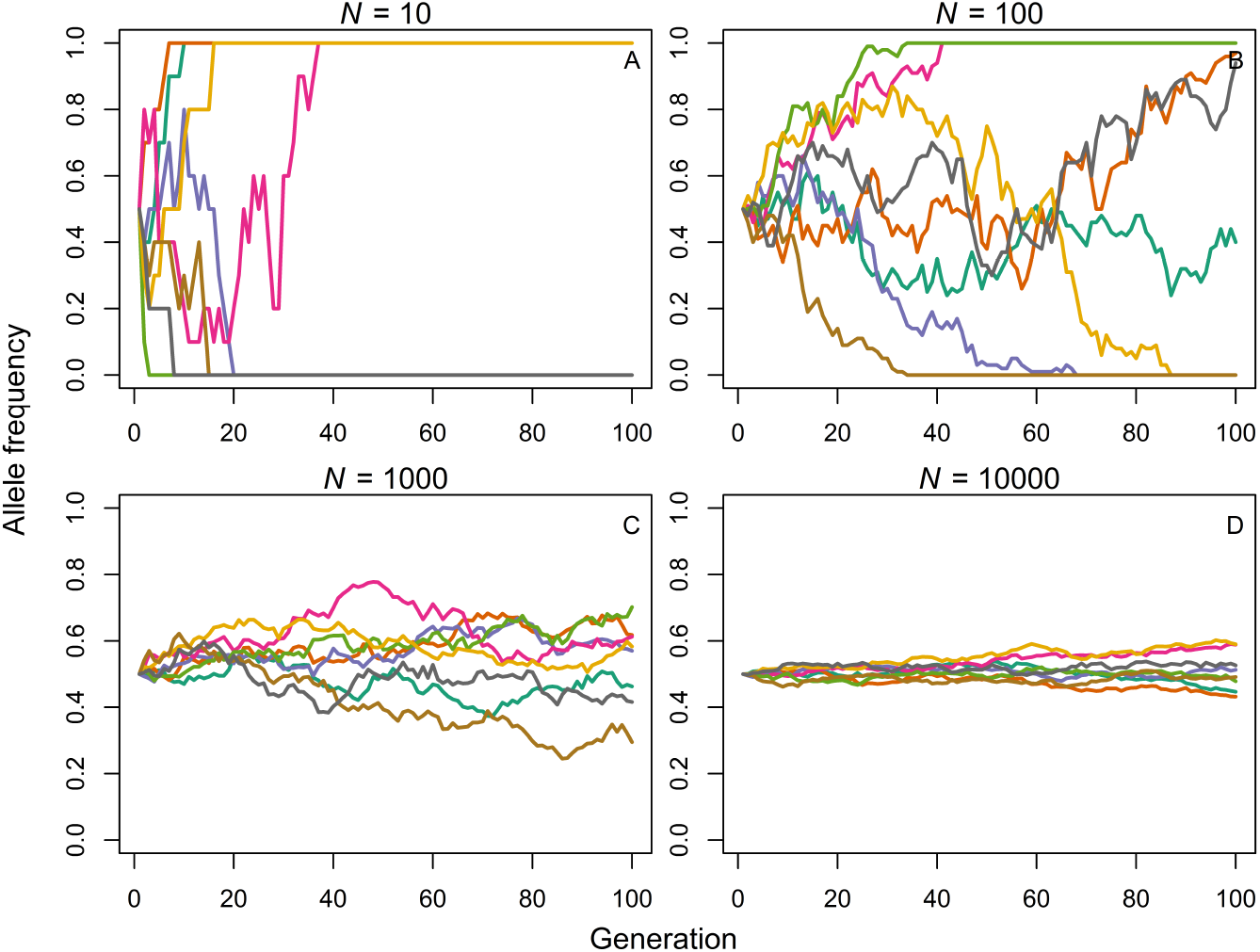
Drift. Frequency of a focal allele over 100 generations. Each line shows evolution in a separate population of size *N* = 10 (A), *N* = 100 (B), *N* = 1000 (C) or *N* = 10000 (D). All simulations start at the same initial allele frequency of 0.5.

### 2.2. Effective population size

The effective size of a population, *N*_*e*_, is a fundamental concept in population genetics that allows researchers to quantify the effect of drift on evolution [Wright, 1931, Kimura and Crow, 1963, Charlesworth, 2009]. *N*_*e*_ is defined as the size of an idealized Wright-Fisher population that is identical in some key measure of genetic drift to a particular study population. Jointly with the mutation rate, *N*_*e*_ determines the expected number of neutral or weakly selected genetic variants maintained in a population and is thus important for correctly calculating the variability in a population. In combination with the strength of selection, *N*_*e*_ also governs how effective selection can be in spreading favourable mutations and eliminating deleterious ones thus shaping the course of adaptive evolution [Charlesworth, 2009].

Different aspects of the evolution of the Wright-Fisher population have been used to define *N*_*e*_. These most often agree but diverge under some circumstances relevant to cultural systems. Therefore, we consider two commonly used measures, inbreeding effective population size, 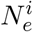, and variance effective population size, 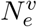. In appendix 3, we provide derivations for haploid populations for situations when (1) there is variation in offspring numbers and (2) population sizes might differ between parental and offspring generation. The identity-by-descent (or inbreeding) effective population size 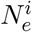 utilizes the fact that, for finite populations, there is a certain probability for two randomly selected individuals to come from the same parent. It is calculated as:

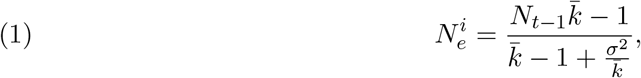

where *N*_*t-*1_ is the census size in the parental generation, 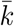 is the mean number of offspring and *s*^2^ is the variance in offspring number among members of the parental generation.

The variance effective population size 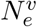, in contrast, focuses on the amount of random variation in allele frequencies from one generation to the next and can be calculated as:

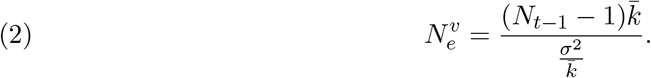

Note that the inbreeding effective number is intimately related to the size of the population in the parental generation, while the variance effective number is related to the size of the population in the offspring generation [Crow and Kimura, 1970]. To understand why, imagine two parents, one carrying allele *A*, one carrying allele *B*, randomly producing a large number of offspring. While this scenario will result in a high probability that two offspring share the same parent, the frequency of both alleles will still be close to 50% in the offspring generation (i.e. low inbreeding effective size and high variance effective size). If, on the other hand, a large number of parents produces only a handful of offspring, there will only be a small probability that two offspring share the same parent, while allele frequencies will differ greatly among generations (i.e. high inbreeding effective size and low variance effective size).

The inbreeding effective number is appropriate when researchers are interested in the change in homozygosity due to random drift. The variance effective number, in contrast, is appropriate when researchers are interested in the amount of gene-frequency drift or the increase in variance among subgroups [Kimura and Crow, 1963, Crow and Denniston, 1988]. Despite these differences, for constant population sizes (i.e. 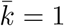), both effective numbers agree and equations simplify to:

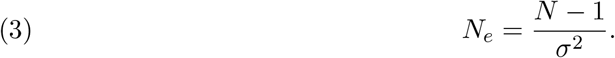

In the following, we will only differentiate between the two formulations when necessary and otherwise use the simplified version given by Eq. (3).

## 3. Determinants of effective population size in cultural evolution

Researchers have identified several factors influencing *N*_*e*_ in genetic evolution [see e.g. Charlesworth, 2009]. In the case of cultural evolution, the relationship between census population size and the effective size may be considerably more complex. For example, the mode of transmission has been shown to be an important factor in determining effective size in genetic systems. In the case of culture, there are many more possible modes of transmission [Cavalli-Sforza and Feldman, 1981, Boyd and Richerson, 1985, Kendal et al., 2018] each of which may have unique effects. Therefore, in order to correctly use the concept of an effective size in cultural systems, a uniquely cultural theory must be developed.

Traits that evolve culturally do so through processes of innovation and cultural transmission. So, we now consider cultural rather than biological reproduction and allow 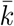 to represent the average number of naive individuals to which a role model transmits their cultural trait. Similarly, *s*^2^ represents the variance in this ‘cultural influence’. Eq. (3) implies that, in general, increasing the variance among individuals in the number of cultural offspring they leave reduces the effective population size, and decreasing that variance increases *N*_*e*_. Thus, it is clear that any process that systematically alters the way in which cultural role models are chosen, through the mode of transmission, demography or social network structure, will change the variance in cultural influence and the effective population size for culturally evolving traits.

### 3.1. Simulation set-up

As the derivation of analytical results for effective population sizes becomes unfeasible for most of the situations considered in this paper, we develop a simulation framework based on the Wright-Fisher dynamic [Kimura and Crow, 1964]. We consider a population of census size *N* where individuals are characterized by the variant of a single cultural trait they have adopted. In each time step, a new generation of individuals is formed and each naive individual adopts its cultural variant, if not specified differently, through unbiased cultural transmission from the previous generation. In more detail, the probability that a naive individual chooses variant *i* of *M* cultural variants present in the previous generation is given by:

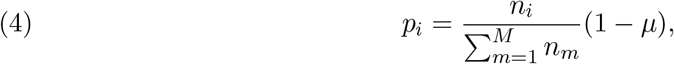

where *n*_*i*_ is the frequency of variant *i* in the appropriate pool of role models. With probability *µ* an innovation takes place and a new, not previously seen variant is introduced into the cultural system. To calculate the effective population size in various scenarios, we first let the system evolve through unbiased transmission until it reaches its equilibrium state. We then run 300 generations assuming the transmission dynamics described below and record the “cultural influence” of each individual in a specific generation by determining its number of cultural offspring in the next generation. This provides us with estimates for 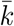 and *s*^2^ (conditioned on the assumed transmission dynamic) needed to calculate effective population sizes according to Eqs. (1) and (2). Additionally, to relate effective population sizes to resulting levels of cultural diversity, we record two diversity measures, the Simpson diversity index (SDI) and the number of unique cultural variants. The SDI is calculated as 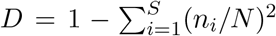, where *n*_*i*_ is the frequency of individuals carrying variant *i* in the population and *S* is the total number of unique cultural variants [Simpson, 1949]. This index ranges from 0 to 1, where high scores indicate high cultural diversity and low scores indicate low cultural diversity.

To account for transmission processes different from unbiased transmission, we adapted Eq. (4). For *one-to-many transmission*, we assign, in each generation, *R* individuals at random as role models and record the frequency *n*_*i*_ in Eq. (4) only from these *R* individuals. For *frequency-dependent transmission*, we assume that the probability for adopting variant *i* of *M* cultural variants present in the population is given by:

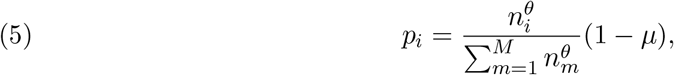

where *n*_*i*_ is the frequency of variant *i* in the whole population and *θ* controls the direction and strength of frequency-dependent bias [McElreath et al., 2008]. When *θ* = 1, cultural transmission is unbiased; as *θ* becomes larger than 1, individuals become increasingly likely to adopt high-frequency variants. When 0 *< θ <* 1, individuals disproportionately adopt low-frequency variants.

To account for *migration* or *cultural exchange*, we use two, initially independent, populations evolving through unbiased transmission. In case of migration, per time step, an average of *mN* randomly chosen individuals from one population permanently migrate to the other population; they carry their cultural variants with them and consequently serve as potential role models for the next generation. The variable *m* controls the migration rate. To keep population sizes constant, the same number of individuals immigrates from the other population. In the case of cultural exchange, per time step, an average of *eN* randomly chosen individuals do not permanently migrate between populations, but are available as additional role models and, thereby, increase the size of the parental generation in both populations. The variable *e* controls the cultural exchange rate.

To account for *social network structure*, we arrange the *N* individuals in the population according to different network topologies (random networks, scale-free networks and small-world networks); this restricts the pool of role models for each individual: the probability of choosing cultural variant *i, p*_*i*_ as given in Eq. (4), is determined only from its direct neighbours in the network. Our aim here is not to replicate realistic social networks but to use prototypical network types to illustrate potential effects of network structure on effective population sizes. Such extreme cases are often useful to identify causal effects and school intuition [see Broom and Voelkl, 2012, Giaimo et al., 2018, for similar analyses for genetic evolution]. All networks considered here are undirected and are generated as follows:

1. Random networks: In random networks, any two individuals have the same probability of being connected. The Erdős-Rényi model generates such a graph by starting with a set of *N* isolated nodes and creating every possible edge with the same constant probability *p* [Erdős and Rényi, 1960]. For undirected graphs, there are 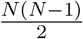 possible ties and *p* gives the expected proportion of those potential ties that are realized in the network (i.e. the network density).
2. Scale-free networks: A network is said to be scale free if the fraction of nodes with degree *k* follows a power law *k*^*-α*^, where *α >* 1. The Barabási-Albert model is an algorithm that uses a preferential attachment mechanism to generate such networks [Barabási and Albert, 1999, Albert and Barabási, 2002]. Here, it is assumed that new nodes are added to the network two at a time. New nodes are connected to existing node *i* (out of *J* total nodes) with a probability *P*_*i*_ that is proportional to the number of links *k*_i_ that a node already has: 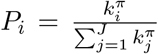. That is, well-connected nodes are likely to get even more connected over time. The power of this preferential attachment is controlled by a parameter *π*, where *π* = 1 produces linear preferential attachment, 0 *< π <* 1 produces “sub-linear” attachment, and *π >* 1 produces “super-linear” attachment.
3. Small-world networks: Small-world networks are graphs with short average path lengths between nodes and a high clustering coefficient. High clustering means that nodes that you are connected to are also likely to be connected to each other (e.g. most of your friends are also friends among themselves). The Watts-Strogatz model creates a small-world network in two basic steps [Watts and Strogatz, 1998]: We start with a lattice of *N* nodes with each node being connected to its *K* closest neighbors on either side. Each edge in the network is then rewired with a certain probability *p*_*r*_ while avoiding duplicates or self-loops. After the first step the graph is a perfect ring lattice. So when *p*_*r*_ = 0, no edges are rewired and the model returns a ring lattice. In contrast, when *p*_*r*_ = 1, all of the edges are rewired and the ring lattice is transformed into a random graph.

All simulation results shown in the following are based on 1000 independent simulations per parameter combination.

### 3.2. Process of cultural transmission

We already know from genetic studies that the mode of inheritance can greatly alter effective population size [e.g. Charlesworth, 2009]. In cultural systems, the ways in which cultural variants can be passed on between individuals, from cultural ‘parents’ to cultural ‘offspring’, are even more numerous and complex [e.g. Cavalli-Sforza and Feldman, 1981, Boyd and Richerson, 1985, Kendal et al., 2018]. In the following, we analyze how processes of cultural transmission can influence effective population size. We consider two transmission processes, *one-to-many transmission* and *frequency-dependent transmission*, whose internal dynamics differ in interesting ways. While the number of transmitting individuals per generation is fixed under one-to-many transmission, it emerges dynamically from the interplay between the transmission mechanism and the frequency spectrum of the cultural variants under frequency-dependent transmission.

#### 3.2.1. One-to-Many Transmission

We define one-to-many transmission as the situation where only a small, pre-defined, number of individuals can transmit their cultural variant to members of the next generation. This transmission process may drastically change the variance in cultural influence and, thus, the effective population size depending on the number of role models, *R*. Here, *R* individuals can pass on their variant and *N - R* individuals cannot. In other words, each generation, we have a transmitting sub-population of size *R* and a non-transmitting sub-population of size *N - R*. In this case, the variance of cultural influence can be calculated as follows (see appendix 4 for the full details):

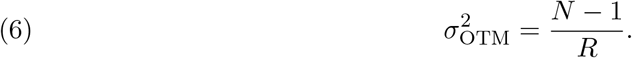

As *R* increases, i.e. as more role models have the chance to pass on their cultural trait, the variance of cultural influence in the population decreases, and for *R* = *N* we recover the expression for the variance of cultural influence in the standard Wright-Fisher model. Thus, the effective population size, *N*_*e*_, for our one-to-many transmission model is simply the size of the transmitting sub-population per generation:

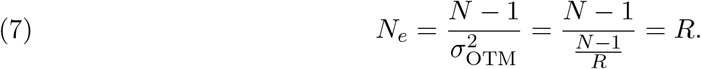

Figs. 2A and B show the variance in cultural influence, given in Eq. (6), and the effective population sizes, given in Eq. (7), for different *R* values. The grey dots represent the mean values of the effective population size generated by the simulation model described in section 3.1 and, reassuringly, analytical and simulation results match very well. When everyone is a potential role model (i.e. *R* = *N*), we recover the standard Wright-Fisher model with a variance of cultural influence of approximately 1 and an effective population size of *N*_*e*_ = *N*. Restricting the pool of role models results in an increased variance in cultural influence and decreased effective population size. In the extreme case where the whole population learns from only one individual per generation, the effective population size is 1.

**Figure 2.**
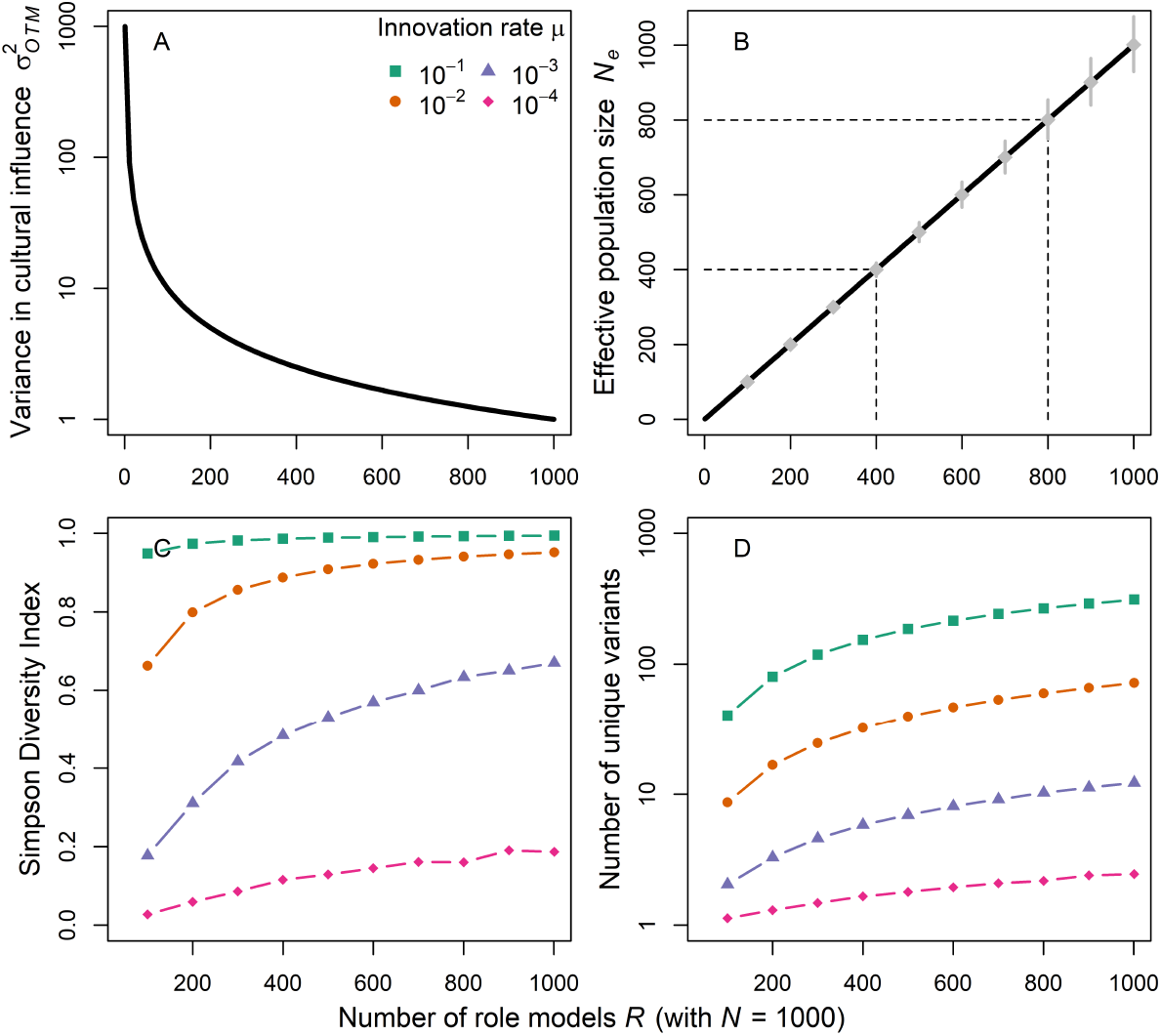
One-to-many transmission. Variance in cultural influence 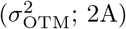, effective population size (*N*_*e*_; 2B), mean Simpson diversity (2C) and mean number of unique cultural variants (2D) for different numbers of cultural role models *R* (with census population size *N* = 1000). Analytical results were confirmed by stochastic simulations. Grey diamonds in plot 2B show means and 90% prediction intervals for 1000 independent simulations (with *µ* = 10^*-*4^).

Figs. 2C and D describe the cultural composition of the population at equilibrium by recording the level of diversity via the Simpson index and number of unique cultural variants in the population. Levels of cultural diversity are jointly determined by the effective population size and innovation rate *µ* (remember that census size is always constant).

#### 3.2.2. Frequency-dependent transmission

We now turn to frequency-dependent cultural transmission where the number of transmitting individuals is not fixed but emerges from the interplay between the transmission process and the frequencies of cultural variants. This form of transmission is well-documented in both human [e.g. Deffner et al., 2020, Van Leeuwen et al., 2018] and non-human animals [e.g. Aplin et al., 2017, Danchin et al., 2018]. Positive frequency-dependent transmission, or conformity, occurs when the most common variants in a population are disproportionately more likely to be adopted. In contrast, negative frequency-dependent transmission, or anti-conformity, occurs when the rarest variants are disproportionately more likely to be copied.

This dynamic is modelled in our simulation framework through Eq. (5). After a burn-in phase under unbiased transmission, i.e. *θ* = 1, we change the *θ* value and record how effective population sizes change over time (see Fig. 3). We start by analysing relatively strong frequency-dependent transmission which results in situations where almost the whole population adopts the same cultural variant (for conformity; *θ* = 1.5, right column) or all cultural variants have similar frequencies (for anti-conformity; *θ* = 0.5, left column). Fig. 3 shows that the change in transmission process leads to an immediate, and partly substantial, decrease in effective population size, followed by a fast recovery. The severity of the peak as well as the new equilibrium after the change is influenced by the innovation rate.

**Figure 3.**
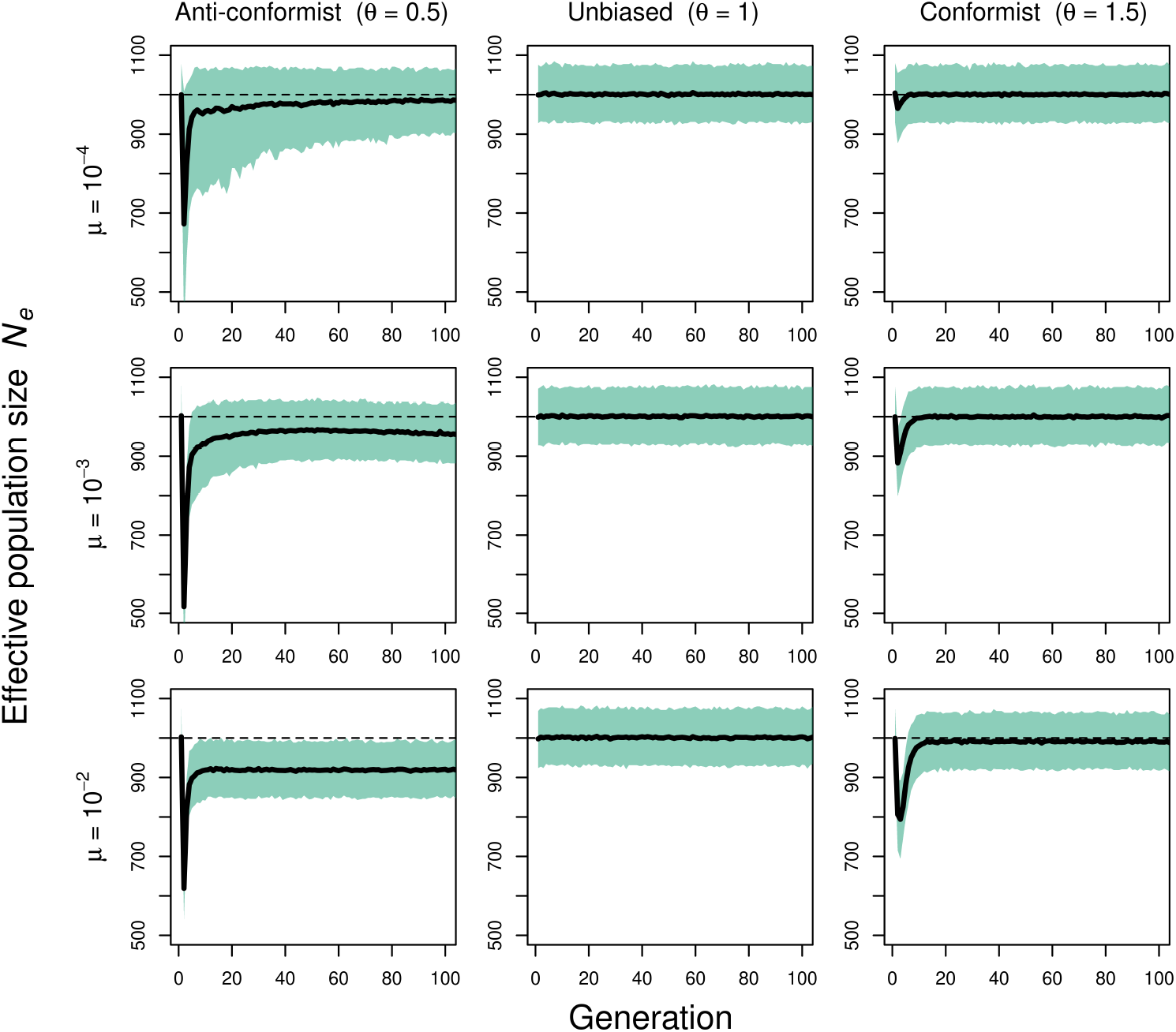
Frequency-dependent transmission. Effective population size (including 90% PIs) for anti-conformist (*θ* = 0.5; left), unbiased (*θ* = 1; center) and conformist transmission (*θ* = 1.5; right) and different innovation rates *µ*. Plots show trajectories for 100 generations after switch in transmission mode (1000 independent simulations; *N* = 1000).

Strong conformity substantially increases the probability that the most common variants are adopted and, therefore, reduces the number of transmitting individuals. This increases the variance in cultural influence and decreases the effective population size. As time progresses, one variant spreads trough the population and almost reaches fixation. Because there is no variation in variant frequency for conformity to act on anymore, every individual (at least every individual that does not carry an innovation) is equally likely to pass on their variant to the next generation; this resembles the situation in the standard Wright-Fisher model. Consequently, at equilibrium *N*_e_ ≈ *N*. We note that in the case of conformity, at equilibrium, higher innovation rates only result in a very slight decrease in effective population size (see Fig. 3 bottom row, right) as innovations are quickly driven to extinction.

Strong anti-conformity substantially increases the probability that the rarest variants are copied, again reducing the number of transmitting individuals. As time progresses, variants become equally distributed and, consequently, individuals do not differ greatly in the likelihood of passing their variants to the next generation. But in contrast to the conformist situation, the effective population size at equilibrium is greatly influenced by the innovation rates; per definition, innovations are rare and, thereby, the target of anti-conformity. The higher the innovation rate, the more likely a variant is present in the population at low frequency and, therefore, the higher the differences between individuals in their cultural influence.

Importantly, we note that the dynamics displayed in Fig. 3 only occur under relatively strong frequency-dependent transmission. In appendix 5, Fig. S1, we show that weaker forms of frequency-dependent transmission leave effective population sizes largely unchanged as now the change in transmission mode does not generate sufficiently large differences in individuals’ likelihood to pass on their cultural variant.

Summarizing, cultural transmission processes different from unbiased transmission do not necessarily lead to a divergence between census and effective population size. This only happens if transmission processes, such as one-to-many and strong frequency-dependent transmission, produce substantial heterogeneity in the probability with which individuals pass on their cultural variant to the next generation, i.e. their cultural influence.

### 3.3. Population connectedness

In the previous sections, we analyzed the impact of different processes of cultural transmission on the effective size of a cultural population. We now turn to the question of how population properties themselves might influence *N*_*e*_. Empirical tests of demographic hypotheses often consider connectedness among groups, which has been assumed to change the effective size of the populations under consideration [Henrich et al., 2016, Derex and Mesoudi, 2020]. We start by analyzing the effects of population connectedness in the form of migration and cultural exchange.

Fig. 4 shows effective population sizes and diversity indices for various degrees of migration on the left and cultural exchange on the right. Irrespective of its rate, migration as we defined it influenced neither inbreeding nor variance effective population size. While introducing new variants into the population, migration in our model does not systematically change the probability individuals get to pass on their cultural traits. As population sizes are constant and individuals still learn from random members of the parental generation, whether they are recent immigrants or not, this scenario corresponds to the standard Wright-Fisher population. Although effective numbers remain unchanged, even small amounts of migration increase both measures of cultural diversity compared to isolated populations. Further raising migration rates leaves diversity largely unchanged indicating that being connected through migration at all has the largest impact on diversity irrespective of the specific rate. Note that if migration increases the census size in a focal population, the effective size tracks this increase (see Fig. S2 in appendix 5 for a scenario where constant immigration raises the effective size and cultural diversity in a focal population).

**Figure 4.**
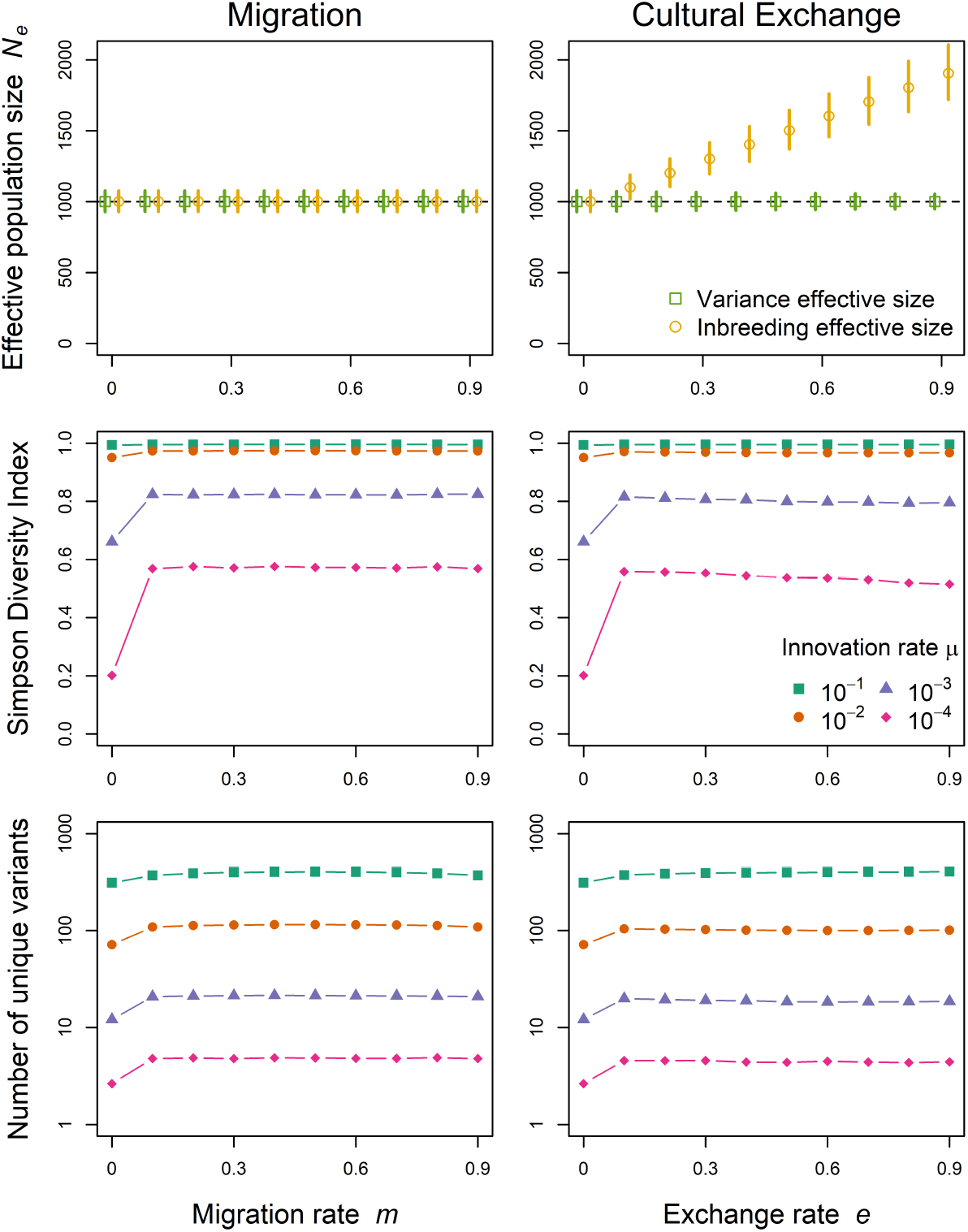
Population connectedness. Variance (green squares) and inbreeding (yellow circles) effective population sizes (including 90% PIs; top), Simpson diversity (center) and mean numbers of unique variants (bottom) for different migration rates *m* on the left and cultural exchange rates *e* on the right. We need to differentiate between effective size formulations because population sizes might differ between parental and offspring generations. Results come from 1000 independent stochastic simulations with census population size *N* = 1000.

Cultural exchange, on the other hand, increases the inbreeding effective population size but not the variance effective size. Why is this? In case of cultural exchange, there are *N* (1 + *e*) individuals in the parental generation that pass on their cultural variants to only *N* individuals in the offspring generation. For the inbreeding effective size, this reduces the probability that two randomly picked individuals copy the same role model in the parental generation as there is now a greater pool of individuals to learn from. In contrast, for the variance effective size, only the number of learners matters such that changing the number of role models has no effect. In practice, this means that in cases where cultural exchange, or similar processes, are important features of a population’s cultural life, the relationship between census population size and effective population size will depend on the measure chosen. Looking at diversity, small rates of cultural exchange lead to the strongest increase in Simpson diversity and the number of unique variants independently of the effective population size.

Summarizing, the effects of connectedness on effective population sizes are subtle, probably difficult to detect, and depend on the exact form of connectedness and effective size formulation.

### 3.4. Social network structure

So far, we have considered panmictic (or “well-mixed”) populations. Real populations, however, are often highly structured in terms of kinship, peer relationships or social class, all influencing who individuals are most likely to interact with and learn from [Knox et al., 2006, Borgatti et al., 2018, Derex and Mesoudi, 2020].

To reflect individual heterogeneity in potential role models and mimic different ways of information flow, we arrange the *N* individuals of the population in networks with different properties (see Fig. 5, top row, and section 3.1 for a more detailed description).

**Figure 5.**
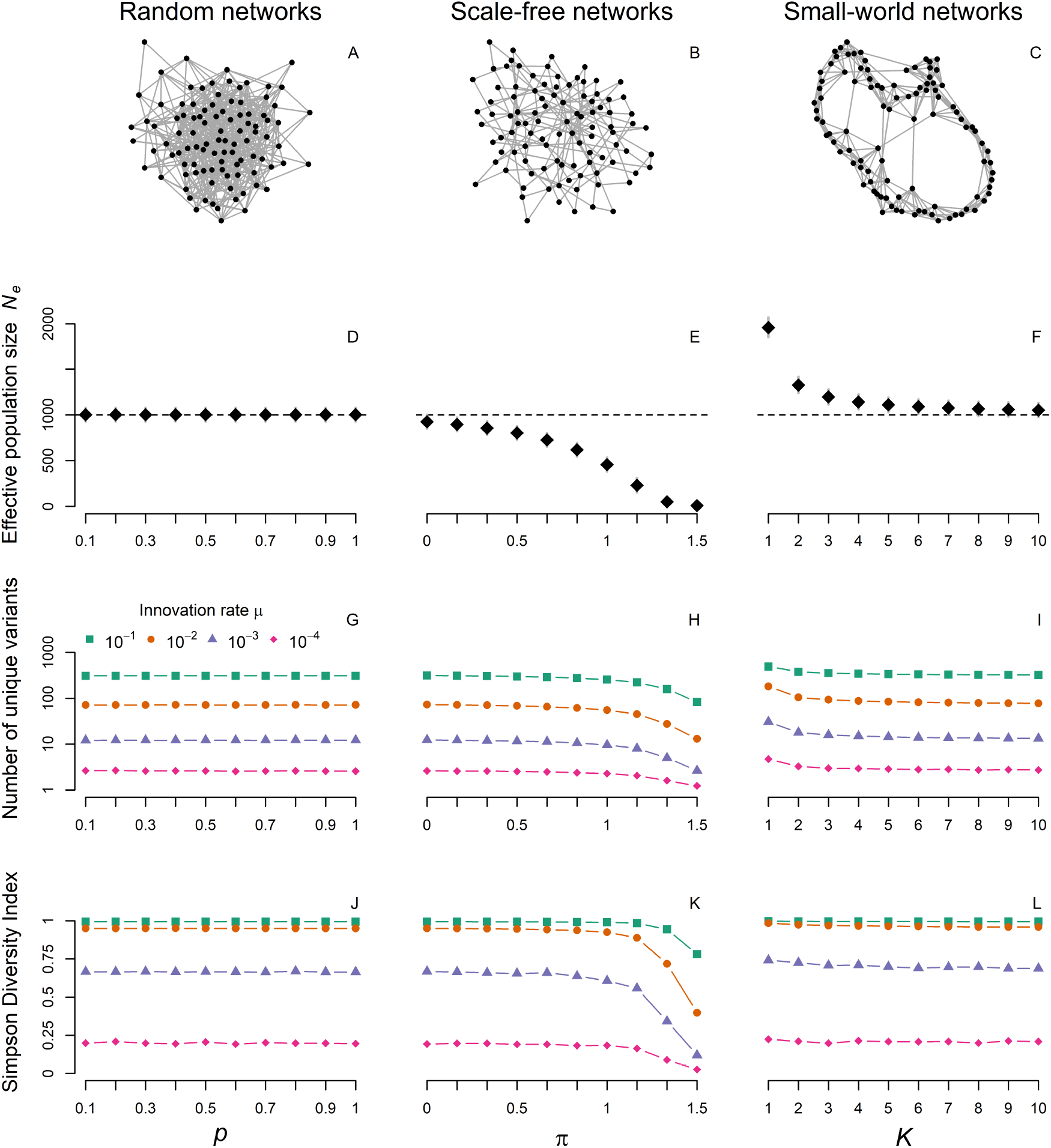
Social network structure. Exemplary networks (A-C), effective population sizes (with 90% PIs; D-F), mean numbers of unique cultural variants (G-I) and Simpson diversity indices (J-L) for random (Erdős-Rényi), scale-free (Barabási-Albert) and small-world (Watts-Strogatz) networks. Parameter *p* gives the probability any two nodes are connected in random networks, *π* is the power of preferential attachment creating scale-free networks and *K* represents the number of initial neighbors on each side in small-world networks (with *p*_*r*_ = 0.01). Note that because of structural differences between network types, the ranges of parameter values on the x-axes are not directly comparable. All graphs are created using the *igraph R* package [Csardi et al., 2006]. Results come from 1000 independent simulations with census population size *N* = 1000. On the top, only 100 nodes are drawn for ease of illustration with *p* = 0.1, *π* = 1 and *K* = 4.

Fig. 5, second row, shows how the way interactions between individuals are structured affects the effective population size. In random networks, irrespective of network density (i.e. the ratio between observed and possible edges), the effective size always equals the census size of the population. As every individual has the same probability *p* of being connected to any other individual, the number of links are binomially distributed and there is no systematic difference in individuals’ probability to pass on their cultural variants. Density also does not affect levels of cultural diversity such that strongly interconnected populations are not more diverse if connections are random.

The situation is very different for scale-free networks. Per definition, their power-law degree distribution implies drastically different levels of cultural influence depending on network position. Individuals central to the network will spread their cultural variant to a great number of individuals while more peripheral individuals will pass on their variant to just a few. This increased variance in cultural influence results in substantially lower effective numbers and also lower levels of cultural diversity. In the extreme case, strong preferential attachment results in a star-shaped network and all individuals will learn from few very central models.

Finally, effective numbers tend to be greater than census numbers for small-world networks. This demonstrates that *N*_*e*_ can also exceed *N* in cultural systems. As a consequence of strongly local cultural transmission, the variance in cultural influence is reduced compared to random or fully-connected networks. That way, rare cultural variants that would otherwise be quickly lost due to drift, can be shared in local clusters maintaining higher levels of cultural diversity. As either the number of initial neighbors, *K*, or the rewiring probability, *p*_*r*_ (not shown here), goes up, we approach a fully connected network where everyone can learn from anyone else and *N*_*e*_ approaches census size *N*.

## 4. Discussion

We have systematically examined effective population size, a concept derived from theoretical population genetics, for culturally evolving traits. The effective size allows us to compare populations, where it would otherwise be difficult to do so. We showed that both modes of cultural transmission and relevant elements of population structure can change the effective size compared to the census size, sometimes considerably. One-to-many and frequency-dependent transmission can substantially lower effective population size with the strongest effects of frequency dependence occurring when the system is out-of-equilibrium. Investigating different forms of connectedness between populations, we found that migration as we define it does not increase *N*_*e*_ and cultural exchange among groups increases inbreeding effective number but not variance effective number. This implies that considerable precision and caution is needed in defining cultural effective sizes. Finally, while random networks with varying densities leave *N*_*e*_ unchanged, scale-free networks tend to decrease and small-world networks tend to increase *N*_*e*_ compared to the census number.

Population size has been invoked to explain several patters of cultural change, most notably the emergence and loss of cultural complexity observed in the ethnographic and archaeological record. Several theoretical models have been developed to better understand the interplay between learning and demography in generating cultural complexity [Shennan, 2001, Henrich, 2004, Powell et al., 2009, Fogarty et al., 2017]. Although quite diverse in terms of underlying mechanisms, these models generally agree in predicting more complex cultural repertoires in larger populations. Both real-world ethnographic and archaeological data as well as controlled lab experiments have been used to test the relationship between population size and cultural complexity. Results with both approaches have been mixed with some studies supporting the hypothesis [e.g. Powell et al., 2009, Kline and Boyd, 2010, Derex et al., 2013, Muthukrishna et al., 2014, Kempe and Mesoudi, 2014] but others not [e.g. Collard et al., 2005,0, Caldwell and Millen, 2010, Fay et al., 2019].

These inconsistent findings do not necessarily refute the theoretical models but might instead reflect a poor correspondence between theory and empirical tests. Here, we have demonstrated that obtaining correct and comparable values for the population size in complex cultural scenarios is not straightforward. In order for the results from theoretical models to apply correctly to empirical cases, we need to ensure that model parameters are correctly translated into measures from complex real-world scenarios.

Our results show that, when there are a few highly influential individuals who—through transmission modes—strongly influence the cultural makeup of the population, the census size and the effective size can diverge. Similarly, where populations are organised into social networks in which individuals are heterogeneous with respect to their degree, the ratio between census and effective size can either increase or decrease depending on network structure. These results also highlight that even relatively small populations might be able to maintain comparatively high levels of cultural diversity if connections are structured in a certain way. Through predominantly local transmission in small-world networks, cultural variants can persist in parts of the network over long periods of time resisting the effects of drift. In this case, larger local clusters or more links between clusters somewhat counter-intuitively reduce the effective population size even though individuals now share ties with more potential cultural models.

In cultural evolutionary studies, effective size has mostly been invoked as a rhetorical device to explain why the demographic hypothesis fails to predict observed levels of complexity in certain cases [however, see Premo, 2016, for a first formal approach]. Our results suggest that this reasoning is too simplistic. For instance, the effects of interconnectedness depend not only on the exact process of exchange but also on the way effective size is defined. Neither migration nor cultural exchange, as we have modelled them, have consistent effects on the effective population size. These results do not imply that connectedness between populations is not an important factor for cultural dynamics, rather that its effect is likely not through increasing the effective size of a population. The finding that small amounts of cultural exchange result in the most diverse populations confirms previous theoretical results suggesting that partial connectivity among populations maximizes cultural accumulation [Baldini, 2015, Derex et al., 2018].

Overall, these results highlight that census numbers cannot generally be relied on when evaluating hypotheses about the effects of demography on culture. It is the effective size that matters and inferring effective population sizes requires detailed knowledge about underlying cultural and demographic processes. To our knowledge, there are no existing methods applicable to estimating effective size in complex cultural systems [for genetic data, see Foll et al., 2015, for an approximate Bayesian computation method to infer genome-wide average effective population size]. The basic theory of *N*_*e*_ in cultural systems is complicated and in need of considerable development before estimation could become feasible through, for instance, generative inference [Kandler and Powell, 2018]. Estimation methods will need to take account of several relevant mechanistic deviations from ideal models simultaneously, where, for example, transmission modes and social networks both play important roles.

Our results can also be used as more informal heuristics to decide when census numbers could be used as proxies for the theoretically relevant effective numbers. In societies where cultural influence is highly skewed in favor of a small elite, for instance, there is no reason to expect that the overall size of that population should be related to cultural complexity. In relatively egalitarian societies, in contrast, where all individuals are equally likely to transmit their ideas and behaviors, census size could be a reasonable approximation of effective size. With respect to transmission biases (or social learning strategies), our results suggest that census numbers might be used as proxies for *N*_*e*_ as long as biases are relatively weak and do not drastically change the relative success of certain cultural variants. Further, if due to recent transformative events, cultural system are out of equilibrium, researchers should not expect census numbers to conform to the theoretically relevant quantity. Finally, if social network structure prevents ideas from flowing freely through the community, census numbers might still be appropriate to test demographic hypotheses as long as connections are relatively random. Most real-world social ties, however, are unlikely to be random and, in that case, census population size might as well under- or overestimate the effective size of cultural populations depending on exact network configurations.

In summary, in order to use the concept of effective population size as an explanatory tool in cultural systems, we must first understand how uniquely cultural processes impact its calculation and use this understanding to develop sophisticated estimation methods capable of capturing the complexity of real world cultural dynamics.

## Data Availability

This manuscript does not contain any empirical data. Simulation and plotting code necessary to reproduce all results and figures in the manuscript can be found on GitHub: https://github.com/DominikDeffner/CulturalEffectivePopulationSize.

## Acknowledgement

We thank members of the Department for Human Behavior, Ecology and Culture at the Max Planck Institute for Evolutionary Anthropology in Leipzig for constructive discussions and criticisms which helped improving this paper.

## Supplementary material for

### Overview

Appendix (1) shows how to calculate effective population size and expected number of cultural traits for the simple example used in section 1.1. in the introduction.

Appendix (2) explains why both mean and variance in offspring number are approximately 1 for in the haploid Wright-Fisher population.

Appendix (3) shows the derivation of both inbreeding and variance effective population sizes as appropriate for cultural evolution.

Appendix (4) derives the appropriate formula for calculating the pooled variance for transmitting and non-transmitting sub-populations that is used to calculate effective population size for one-to-many transmission.

Appendix (5) contains additional results.

#### 1. Calculations for example used in section 1.1. in the introduction

Remember that we assumed there are two populations, A and B, with current census population sizes of 1000 and 500 individuals, respectively. Population A had a population bottleneck ten generations ago when its census size fell to just 10 individuals before immediately returning to its current size of 1000. If population size is fluctuating over time and we have information from a total of *T* non-overlapping generations, the effective size *N*_*e*_ is given by the harmonic mean (i.e. the reciprocal of the arithmetic mean of the reciprocals) of the population sizes at each point in time *t*:

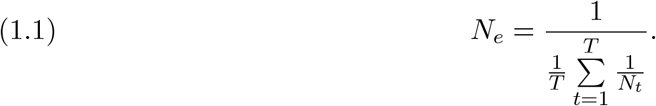

In our example, for population A, *N*_1_ = 10 and *N*_[2:10]_ = 1000, for population B, *N*_[1:10]_ = 500. Plugging these values into Eq. (1.1) gives the effective population sizes of *N*_*e*_ = 91.7 for population A and *N*_*e*_ = 500 for population B.

Based on these effective sizes and assuming a certain innovation rate (here *µ* = 0.1) and unbiased cultural transmission, we can use results from population genetics [see Ewens, 2012, p. 115] to approximate the mean number of cultural traits we expect to see each generation, *E*(*K*), as follows:

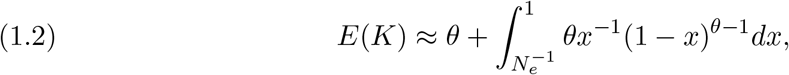

where *θ* = 2*N*_*e*_*µ* is the standard population mutation parameter. For population B with *N*_*e*_ = 500, the expected number of traits is, thus, *E*(*K*) ≈ 223. For population A with *N*_*e*_ = 91.7, we expect to see on average *E*(*K*) ≈ 41 traits in a given generation.

#### 2. Properties of the haploid Wright-Fisher population

Both mean and variance in offspring number are roughly 1 for a haploid Wright-Fisher population. Why is this the case? For the Wright-Fisher population with a population size of *N, N* parents are chosen at random with replacement from the full population. From the point of view of any focal individual, the probability of being chosen as a parent to any one of the *N* offspring is *p* = 1*/N*, the probability of not being chosen, correspondingly, is *q* = 1 − 1*/N*. We conduct *N* trials and so the number of offspring produced by any member of the population is a binomially distributed random variable. The mean of this is simply 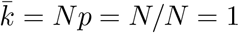. The variance is *σ*^2^ = *Npq*. This is 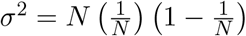. Simplifying, we get 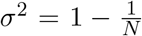. It is common here to neglect terms on the order of 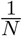; they are very small because *N* is usually very large, giving us *σ*^2^ ≈ 1.

#### 3. Derivation of inbreeding and variance *N*_*e*_

Different aspects of the evolution of the Wright-Fisher population have been used to define *N*_*e*_. We seek expressions for both inbreeding and variance effective numbers for situations where (1) there is variation in offspring numbers and (2) population sizes might differ between parental and offspring generation. Here, we follow the general presentation of Kimura and Crow [1963], but make adjustments for haploid populations where necessary. Haploids are characterized by only one set of variants, whereas diploids are characterized by two sets. In cultural transmission, learners adopt a single variant of a cultural trait from one or several role models, there is no inheritance of two corresponding alleles from sexually reproducing parents.

Consider a population of *N*_*t*−1_ individuals each contributing a variable number, *k*_*i*_, of offspring to the next generation. In general, the mean number of offspring is

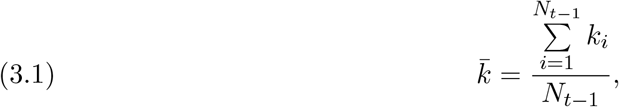

and the variance in offspring number is

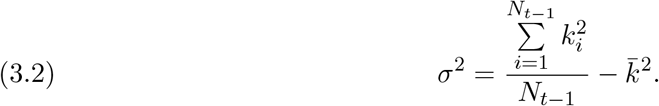

##### 3.1. Inbreeding effective population size

The identity-by-descent (or inbreeding) effective population size 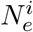 utilizes the fact that in finite populations there is a certain probability that two randomly selected individuals in generation *t* are descendant from the same parent. As this single-generation probability of identity by descent, *P*_*t*_, in the ideal Wright-Fisher population is simply 1*/N*_*t*−1_, we can use an estimate of this probability in the real population to calculate the effective population size as 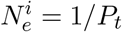. For our example, the number of ways in which two offspring from a given parent *i* can be selected is *k*_*i*_(*k*_*i*_ − 1)*/*2. Summing over all members of the parental generation, the total number of ways in which two offspring from the same parent can be selected is 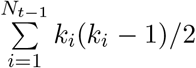, whereas the total number of offspring pairs is 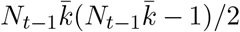. Dividing the former by the latter, *P*_*t*_ can thus be calculated as

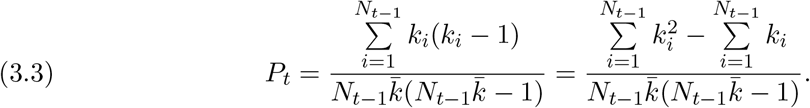

From the definition of the mean (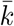; see equation 3.1),

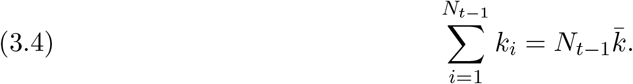

and from the definition of the variance (*σ*^2^; see equation 3.2),

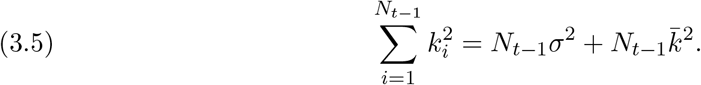

Substituting these into equation 3.3 results in

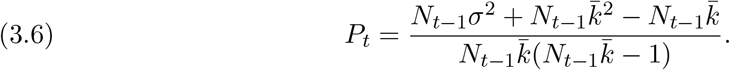

Rearranging, we can calculate the inbreeding effective number for haploid populations as

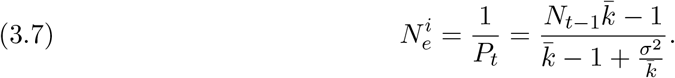

#### 3.2. Variance effective population size

The variance effective population size 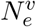, in contrast, focuses on the amount of random variation in allele frequencies from one generation to the next. Assume that *p* is the frequency of an allele in an ideal Wright-Fisher population of size *N*. The sampling variance of the gene frequency drift from parent to offspring generation is *V*_*δp*_ = *p*(1 − *p*)*/N* and, therefore, 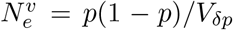. Similarly to above, we seek an expression for *V*_*δp*_ to infer the corresponding effective number. Again, assume there is a population of *N*_*t*−1_ individuals each contributing a number of *k*_*i*_ offspring to the next generation. Let *p* denote the frequency of allele *A* in the population and let *n*_1_ be the number of individuals carrying allele *A* (*n*_1_ = *N*_*t*−1*p*_). The number of *A* alleles contributed to the next generation is 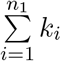. Thus, the increment in *A* alleles from generation *t* − 1 to generation *t* is given by:

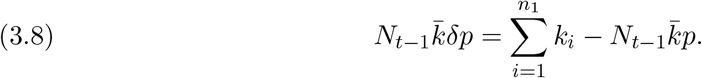

Expressing *p* as 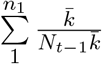 and substituting into equation (3.8) gives,

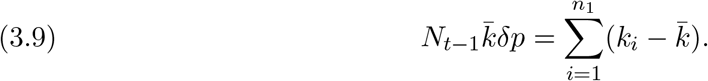

Using expectation notation, we now write an expression for the variance in the change in *p, V*_*δp*_ = **E**[*δp*]^2^.

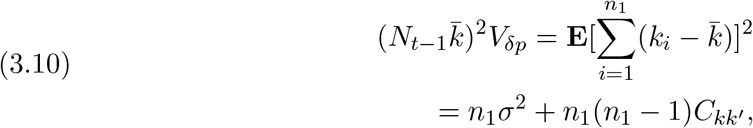

where 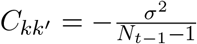 is the covariance in offspring number for randomly selected pairs from the parental generation. Substituting this and simplifying results in

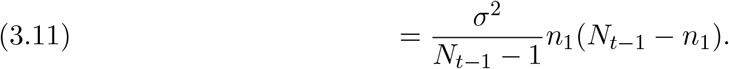

If we then replace *n*_1_ by *N*_*t*−1_*p*, we get

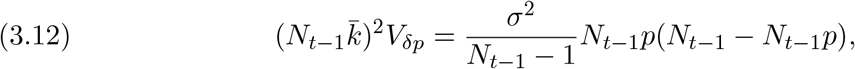

and

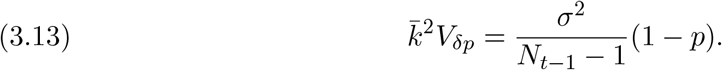

Replacing *V*_*δp*_ with 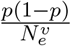 gives

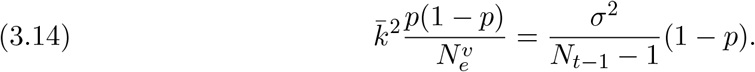

Solving for 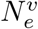 results in

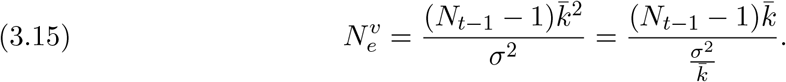

#### 4. Expressing the variance of the full population in terms of sub-population variances and means

The combined variance, 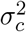, is given by

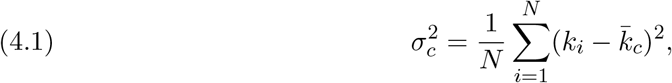

where, again, *N* is the total population size, *k*_*i*_ is the number of cultural offspring of the *i*^*th*^ individual in the parental generation, and 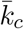 is the combined mean offspring number for the whole population (including both the group of cultural transmitters and the group of non-transmitters). To write a general expression for the variance in the full population, we split the population into two arbitrary groups of size *n*_1_ and *n*_2_ where *n*_1_ + *n*_2_ = *N*. We label the mean offspring number within those subpopulations as 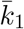 and 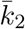 respectively, and the variances in offspring number, similarly, as 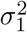 and 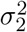.

Now, it is possible to express the variance of the complete population 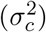 in terms of the means and variances of the two subpopulations.

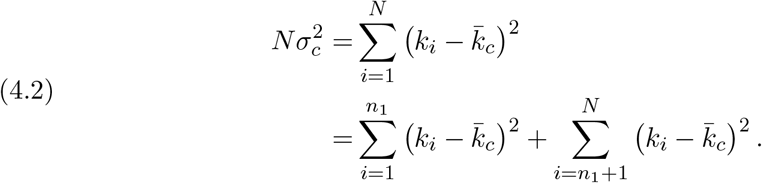

With some rearranging, we get

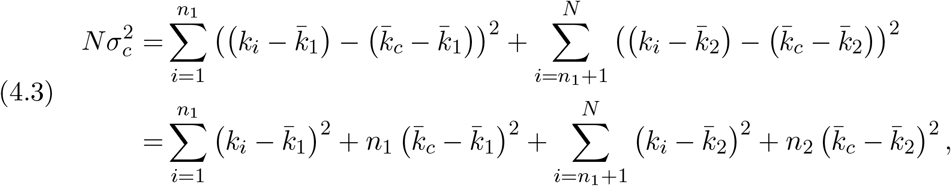

using the fact that in general 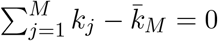. So, we have

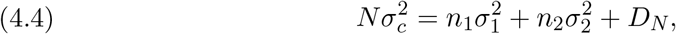

where

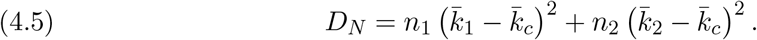

To allow some simplifications we can rewrite *D*_*N*_ as

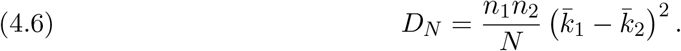

Details of this simplification can be found in O’Neill [2014]. The decomposed expression for 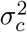, then is

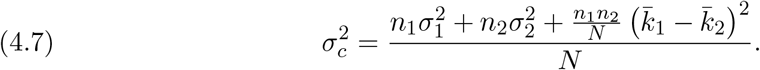

Now, to apply this to our population and to the calculation of effective population size, we make the following assumptions. Population 1 is the transmitting population. Therefore, *n*_1_ = *R, n*_2_ = (*N* − *R*). 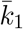 is the mean number of cultural offspring per individual in the transmitting sub-population. Given that there are *N* learners and *R* role models, we get that 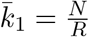, and the mean for the other sub-population is 0, since they cannot pass on their trait 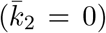. The variance for the non-transmitters 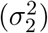 is the same, 0. The variance for the transmitting population is obtained in the same way as the variance for the full Wright-Fisher population and is 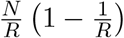.

Inserting these values into equation 4.7 results in

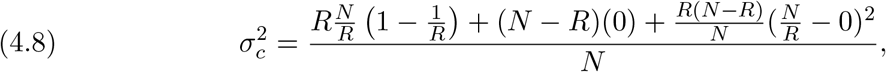

which simplifies to

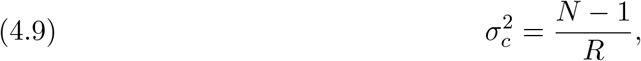

the solution for 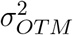 shown in the main text (equation 6).

#### 5. Additional results

**Figure S1.**
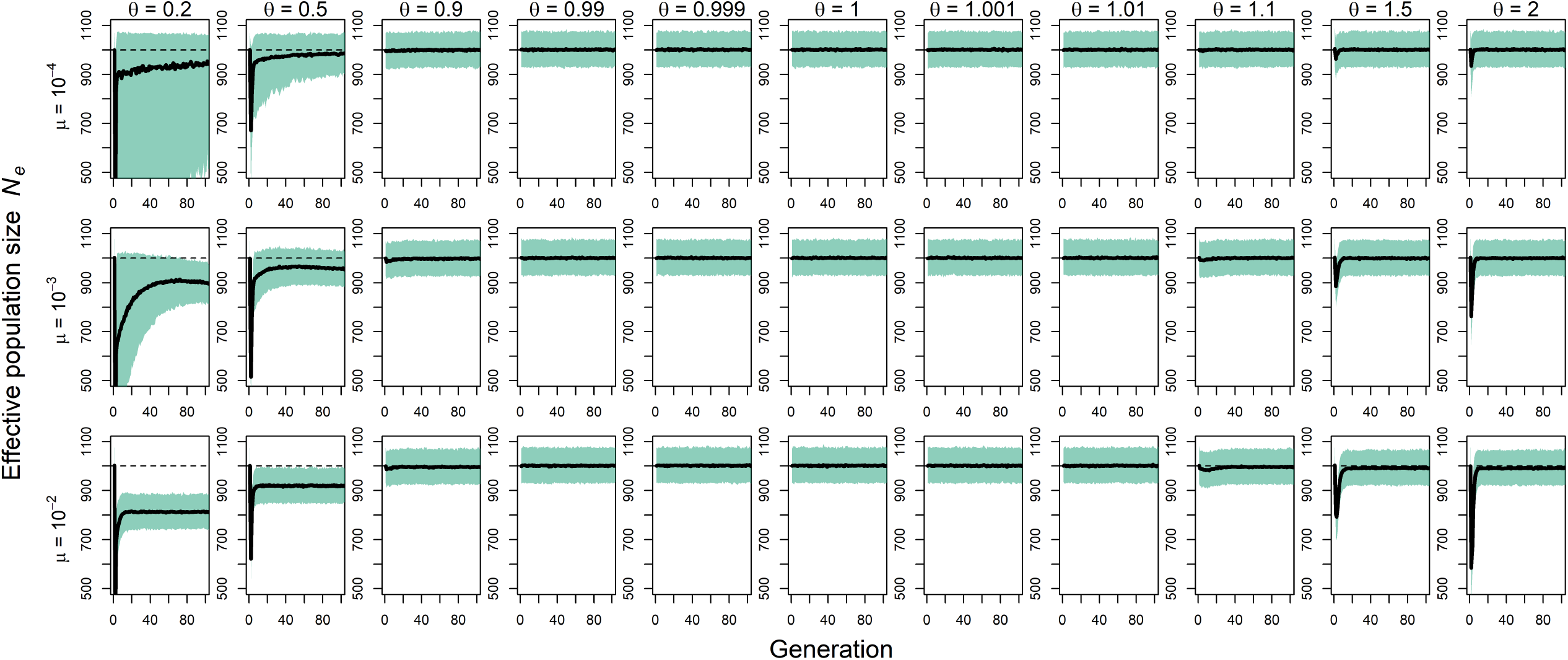
Frequency-dependent transmission for wider ranges of conformity exponent *θ*. Effective population size (including 90% PIs) for different levels of anti-conformist (*θ <* 1), unbiased (*θ* = 1) and conformist transmission (*θ >* 1) and different innovation rates *µ*. Plots show trajectories for 100 generations after switch in transmission mode (1000 independent simulations; *N* = 1000).

**Figure S2.**
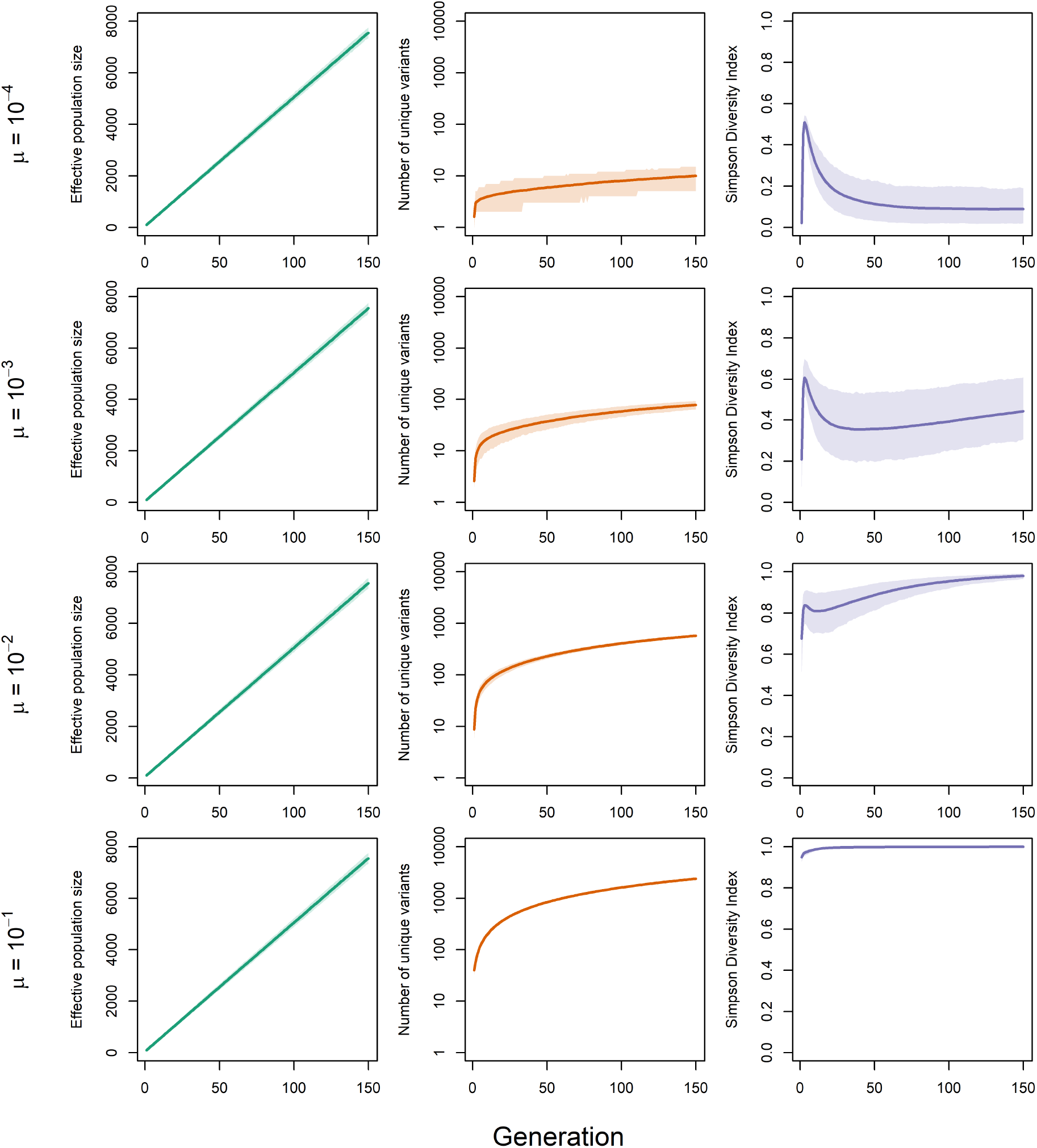
Alternative migration mechanism that gradually increases the census size *N* in a focal population. We start by letting a large source population with *N* = 10000 and a small focal population with *N* = 100 evolve separately until they reach equilibrium; each generation, we then let a fixed number of individuals migrate from the source population to the focal population and record effective numbers and diversity indices in the focal population. Effective population size (left), number of unique cultural variants (center) and Simpson Diversity (right) for different innovation rates *µ*. Plots show trajectories for 150 generations after immigration starts (1000 independent simulations with 50 immigrants per generation).

## References

Luigi Luca Cavalli-Sforza and Marcus W Feldman. Cultural transmission and evolution: a quantitative approach. Princeton University Press, 1981.

Robert Boyd and Peter J Richerson. Culture and the evolutionary process. University of Chicago press, 1985.

Maxime Derex and Alex Mesoudi. Cumulative cultural evolution within evolving population structures. Trends in Cognitive Sciences, 2020.

Sarah Saxton Strassberg and Nicole Creanza. Cultural evolution and prehistoric demography. Philosophical Transactions of the Royal Society B, 376(1816):20190713, 2021.

Warren J Ewens. Mathematical population genetics 1: theoretical introduction, volume 27. Springer Science & Business Media, 2012.

Ronald A Fisher. On the dominance ratio. Proceedings of the Royal Society of Edinburgh, 42:321–341, 1923.

Sewall Wright. Evolution in mendelian populations. Genetics, 16(2):97–159, 1931.

Stephen Shennan. Demography and cultural innovation: a model and its implications for the emergence of modern human culture. Cambridge Archaeological Journal, 11(1):5–16, 2001.

Joseph Henrich. Demography and cultural evolution: how adaptive cultural processes can produce maladaptive losses. the tasmanian case. American Antiquity, 69(2):197–214, 2004.

Adam Powell, Stephen Shennan, and Mark G Thomas. Late pleistocene demography and the appearance of modern human behavior. Science, 324(5932):1298–1301, 2009.

Laurel Fogarty, Joe Yuichiro Wakano, Marcus W Feldman, and Kenichi Aoki. The driving forces of cultural complexity. Human Nature, 28(1):39–52, 2017.

Wendell H Oswalt. An anthropological analysis of food-getting technology. John Wiley & Sons, New York, 1976.

Laurel Fogarty and Anne Kandler. The fundamentals of cultural adaptation: implications for human adaptation. Scientific Reports, 10(1):1–11, 2020.

Robin S Waples, Gordon Luikart, James R Faulkner, and David A Tallmon. Simple life-history traits explain key effective population size ratios across diverse taxa. Proceedings of the Royal Society B: Biological Sciences, 280(1768):20131339, 2013.

Albert Tenesa, Pau Navarro, Ben J Hayes, David L Duffy, Geraldine M Clarke, Mike E Goddard, and Peter M Visscher. Recent human effective population size estimated from linkage disequilibrium. Genome Research, 17(4):520–526, 2007.

Joseph Henrich, Robert Boyd, Maxime Derex, Michelle A Kline, Alex Mesoudi, Michael Muthukrishna, Adam T Powell, Stephen J Shennan, and Mark G Thomas. Understanding cumulative cultural evolution. Proceedings of the National Academy of Sciences, 113(44): E6724–E6725, 2016.

Lukas S Premo. Effective population size and the effects of demography on cultural diversity and technological complexity. American Antiquity, 81(4):605–622, 2016.

Ronald A Fisher. The distribution of gene ratios for rare mutations. Proceedings of the Royal Society of Edinburgh, 50:204–219, 1931.

Sewall Wright. The evolution of dominance. The American Naturalist, 63(689):556–561, 1929.

Joanna Masel. Genetic drift. Current Biology, 21(20):R837–R838, 2011.

Michael C Whitlock and Patrick C Phillips. Drift: Introduction. eLS, 2014.

Motoo Kimura and James F Crow. The measurement of effective population number. Evolution, pages 279–288, 1963.

Brian Charlesworth. Effective population size and patterns of molecular evolution and variation. Nature Reviews Genetics, 10(3):195–205, 2009.

James F Crow and Motoo Kimura. An introduction to population genetics theory. Harper & Row, New York, 1970.

James F Crow and Carter Denniston. Inbreeding and variance effective population numbers. Evolution, 42(3):482–495, 1988.

Rachel L Kendal, Neeltje J Boogert, Luke Rendell, Kevin N Laland, Mike Webster, and Patricia L Jones. Social learning strategies: Bridge-building between fields. Trends in Cognitive Sciences, 22(7):651–665, 2018.

Motoo Kimura and James F Crow. The number of alleles that can be maintained in a finite population. Genetics, 49(4):725, 1964.

Edward H Simpson. Measurement of diversity. Nature, 163(4148):688–688, 1949.

Richard McElreath, Adrian V Bell, Charles Efferson, Mark Lubell, Peter J Richerson, and Timothy Waring. Beyond existence and aiming outside the laboratory: estimating frequency-dependent and pay-off-biased social learning strategies. Philosophical Transactions of the Royal Society B: Biological Sciences, 363(1509):3515–3528, 2008.

Mark Broom and Bernhard Voelkl. Two measures of effective population size for graphs. Evolution, 66(5):1613–1623, 2012.

Stefano Giaimo, Jordi Arranz, and Arne Traulsen. Invasion and effective size of graph-structured populations. PLoS Computational Biology, 14(11):e1006559, 2018.

Paul Erdős and Alfréd Rényi. On the evolution of random graphs. Publ. Math. Inst. Hung. Acad. Sci, 5(1):17–60, 1960.

Albert-Lázló Barabási and Réka Albert. Emergence of scaling in random networks. Science, 286(5439):509–512, 1999.

Réka Albert and Albert-László Barabási. Statistical mechanics of complex networks. Reviews of Modern Physics, 74(1):47, 2002.

Duncan J Watts and Steven H Strogatz. Collective dynamics of ‘small-world’ networks. Nature, 393(6684):440–442, 1998.

Dominik Deffner, Vivien Kleinow, and Richard McElreath. Dynamic social learning in temporally and spatially variable environments. Royal Society Open Science, 7(12):200734, 2020.

Edwin JC Van Leeuwen, Emma Cohen, Emma Collier-Baker, Christian J Rapold, Marie Schäfer, Sebastian Schütte, and Daniel BM Haun. The development of human social learning across seven societies. Nature Communications, 9(1):1–7, 2018.

Lucy M Aplin, Ben C Sheldon, and Richard McElreath. Conformity does not perpetuate suboptimal traditions in a wild population of songbirds. Proceedings of the National Academy of Sciences, 114(30):7830–7837, 2017.

Etienne Danchin, Sabine Nöbel, Arnaud Pocheville, Anne-Cecile Dagaeff, Léa Demay, Mathilde Alphand, Sarah Ranty-Roby, Lara Van Renssen, Magdalena Monier, Eva Gazagne, et al. Cultural flies: Conformist social learning in fruitflies predicts long-lasting mate-choice traditions. Science, 362(6418):1025–1030, 2018.

Hannah Knox, Mike Savage, and Penny Harvey. Social networks and the study of relations: networks as method, metaphor and form. Economy and Society, 35(1):113–140, 2006.

Stephen P Borgatti, Martin G Everett, and Jeffrey C Johnson. Analyzing social networks. Sage, 2018.

Gabor Csardi, Tamas Nepusz, et al. The igraph software package for complex network research. InterJournal, complex systems, 1695(5):1–9, 2006.

Michelle A Kline and Robert Boyd. Population size predicts technological complexity in oceania. Proceedings of the Royal Society B: Biological Sciences, 277(1693):2559–2564, 2010.

Maxime Derex, Marie-Pauline Beugin, Bernard Godelle, and Michel Raymond. Experimental evidence for the influence of group size on cultural complexity. Nature, 503(7476): 389–391, 2013.

Michael Muthukrishna, Ben W Shulman, Vlad Vasilescu, and Joseph Henrich. Sociality influences cultural complexity. Proceedings of the Royal Society B: Biological Sciences, 281(1774):20132511, 2014.

Marius Kempe and Alex Mesoudi. An experimental demonstration of the effect of group size on cultural accumulation. Evolution and Human Behavior, 35(4):285–290, 2014.

Mark Collard, Michael Kemery, and Samantha Banks. Causes of toolkit variation among hunter-gatherers: a test of four competing hypotheses. Canadian Journal of Archaeology/Journal Canadien d’Archéologie, pages 1–19, 2005.

Mark Collard, Briggs Buchanan, and Michael J O’Brien. Population size as an explanation for patterns in the paleolithic archaeological record: more caution is needed. Current Anthropology, 54(S8):S388–S396, 2013.

Christine A Caldwell and Ailsa E Millen. Human cumulative culture in the laboratory: effects of (micro) population size. Learning & Behavior, 38(3):310–318, 2010.

Nicolas Fay, Naomi De Kleine, Bradley Walker, and Christine A Caldwell. Increasing population size can inhibit cumulative cultural evolution. Proceedings of the National Academy of Sciences, 116(14):6726–6731, 2019.

Ryan Baldini. Revisiting the effect of population size on cumulative cultural evolution. Journal of Cognition and Culture, 15(3-4):320–336, 2015.

Maxime Derex, Charles Perreault, and Robert Boyd. Divide and conquer: intermediate levels of population fragmentation maximize cultural accumulation. Philosophical Transactions of the Royal Society B: Biological Sciences, 373(1743):20170062, 2018.

Matthieu Foll, Hyunjin Shim, and Jeffrey D Jensen. Wfabc: a wright–fisher abc-based approach for inferring effective population sizes and selection coefficients from time-sampled data. Molecular Ecology Resources, 15(1):87–98, 2015.

Anne Kandler and Adam Powell. Generative inference for cultural evolution. Philosophical Transactions of the Royal Society B: Biological Sciences, 373(1743):20170056, 2018.

## References

B. O’Neill. Some Useful Moment Results in Sampling Problems. American Statistician, 68 (4):282–296, 2014. ISSN 15372731. 10.1080/00031305.2014.966589.

